# Procedure-Matched Comparison of TNBS/Ethanol Enema and DSS Free-Drinking Models of Ulcerative Colitis in Rats: Immune Endotype Profiling and a Translational Model-Selection Framework

**DOI:** 10.64898/2026.06.03.729947

**Authors:** Yongkang An, Fang Wu, Linhan Zhang, Menghua Shi, Shuangxi Zhang, Xiang’an Zhang

**Affiliations:** Proctology Department, the First Affiliated Hospital of Henan University of Chinese Medicine, Zhengzhou, Henan 450000, China; Acupuncture Department, the First Affiliated Hospital of Henan University of Chinese Medicine, Zhengzhou, Henan 450000, China

**Keywords:** Ulcerative colitis, TNBS, DSS, animal model, immune endotype

## Abstract

**Objective:** Choosing between TNBS/ethanol enema and DSS free-drinking water protocols for establishing rat ulcerative colitis(UC) models remains a practical challenge in preclinical research. Beyond confirming that both methods induce colitis, no prior study has quantified the independent contribution of enema procedural trauma to inflammatory readouts, reported sex-stratified cytokine responses, or formally linked model-specific immune profiles to human UC endotypes.

**Methods:** Ten SPF-grade SD rats(half male and half female) were allocated to a normal control, a procedure-matched saline enema control(anesthesia and enema without chemical agent), a TNBS group(100 mg/kg TNBS + 50% ethanol enema), and a DSS group(3% DSS free-drinking, 7 days). Endpoint assessments included body weight, colon length, macroscopic and histological scoring(modified Macpherson criteria, 0–10), and serum TNF-α and IL-6 by ELISA. Post-hoc exploratory analyses included sex-stratified cytokine sub-analysis and coefficient of variation(CV%) for each readout.

**Results:** Both model groups successfully induced UC(*P<*0.05 vs. controls for all endpoints). Critically, the procedure-matched saline enema control confirmed that colonic inflammation in the TNBS group was attributable to the chemical agent rather than mechanical trauma. TNBS induced significantly higher TNF-α(90.2±10.5 vs. 73.5±9.8 pg/mL, *P<*0.05) and IL-6(66.5±8.3 vs. 53.2±7.6 pg/mL, *P<*0.05) than DSS,consistent with Th1-polarized vs. innate barrier-disruption immune endotypes, respectively. Coefficient of variation analysis indicated greater intra-group variability in TNBS versus DSS, reflecting the procedural dependency of enema-based models.

**Conclusion:** A procedure-matched control design is essential for valid TNBS-vs.-DSS comparisons and has been systematically underutilized in prior studies. Based on immune endotype alignment, TNBS is recommended for screening T-cell-targeted therapies and acute anti-inflammatory agents, while DSS is recommended for epithelial barrier repair and microbiome-modulating interventions. A model-selection decision framework derived from these data is provided to guide preclinical study design.

## 1 Introduction

Ulcerative colitis(UC)^[1-3]^ is a chronic intestinal disease characterized by continuous, nonspecific inflammation of the colonic mucosa. Its incidence is increasing globally, with clinical manifestations including recurrent diarrhea, bloody and mucous stools, and abdominal pain^[4]^. Severe cases can progress to colon cancer, placing a heavy burden on patients and society^[5, 6]^. The etiology and pathogenesis of UC are not fully understood, but are likely related to multiple factors such as genetic susceptibility, gut microbiota dysbiosis, and immune dysfunction^[7-9]^. Animal models are core tools for elucidating the pathological mechanisms of UC and screening potential therapeutic drug[10-12]. TNBS enema and DSS free drinking water method are currently the two most widely used modeling techniques, but they differ in their modeling mechanisms, pathological characteristics, and applicable scenarios, lacking a unified standard for model selection in clinical research^[13-15]^. This study systematically compares the changes in general condition, intestinal morphology, pathology, and serum inflammatory factors in rats induced by the two modeling methods, clarifying the advantages, disadvantages, and applicable scope of each model. This provides a scientific and reliable basis for constructing animal models for basic UC research, improving the reproducibility and specificity of experimental studies.

## 2 Research subjects

Ten SPF-grade SD rats(half male and half female) provided by Liaoning Changsheng Biotechnology Co., Ltd. were selected. Production license number: SCXK(Liaoning)2020-0001, Animal Quality Certificate number: NO. 210726231101752781. All animal procedures were approved by the Animal Ethics Committee of Henan Academy of Chinese Medicine(Ethics approval No. 20230911) and were performed in strict accordance with the ARRIE guidelines^[16]^ and the AVMA Guidelines for the Euthanasia of Animals(2020 edition)^[17]^. Every effort was made to minimize animal suffering and reduce the number of animals used(Figure 1).

**Figure 1:**
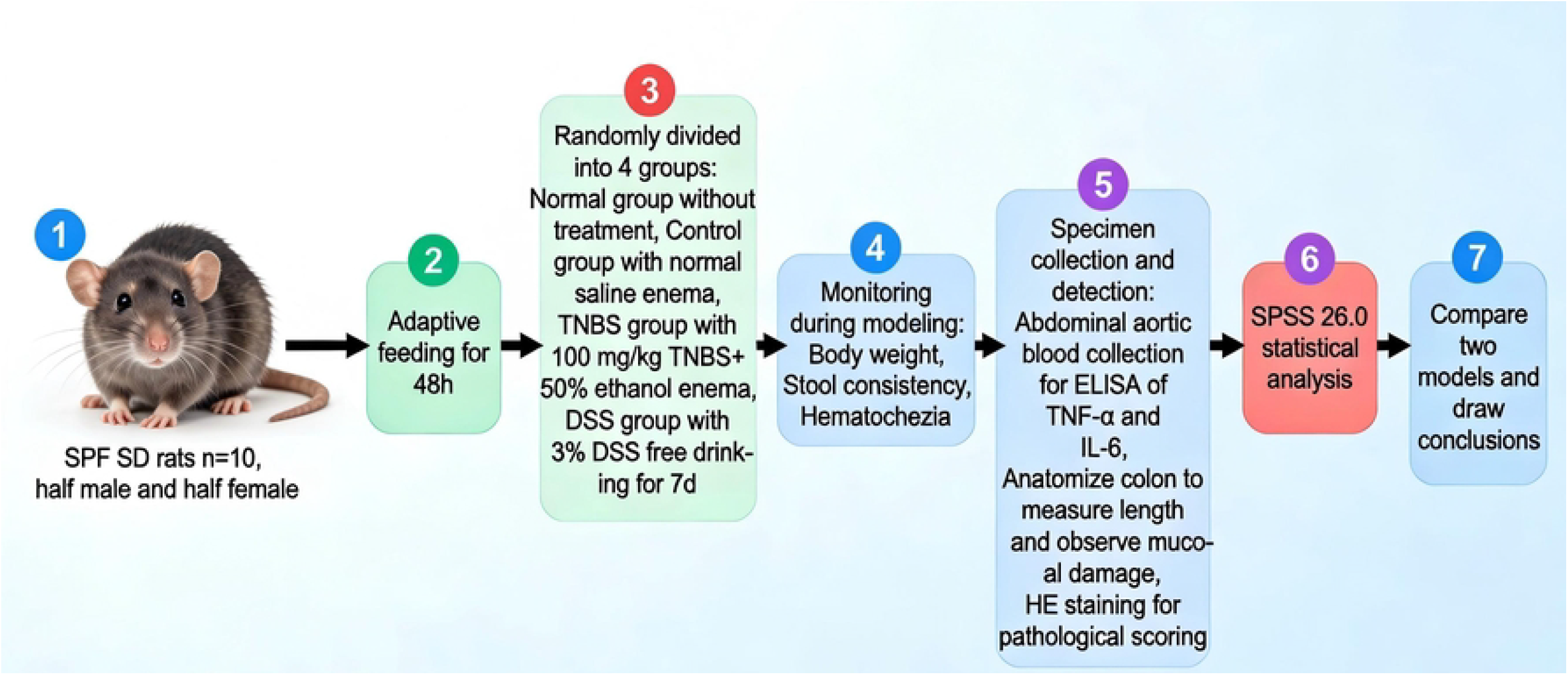
Animal Experiment Flowchart

## 3 Materials

3% sodium dextran sulfate(DSS, molecular weight 36000-50000 Da, Sigma-Aldrich, USA), 2,4,6-trinitrobenzenesulfonic acid(TNBS, purity≥98%, Solarbio Beijing Company), anhydrous ethanol(analytical grade, Sinopharm Chemical Reagent Co., Ltd.), physiological saline, 4% paraformaldehyde fixative(Wuhan Google Biotechnology Co., Ltd.), isoflurane(R510-22-10, Shenzhen Ruiwode Life Technology Co., Ltd.); TNF-α and IL-6 ELISA detection kits(Shanghai Enzyme-Linked Biotechnology Co., Ltd.); disposable blood collection needles, 10ml intravenous infusion set, 5ml centrifuge tubes, 2ml cryovials, rat-specific gavage apparatus(Jiangsu Kangjie Medical Technology Co., Ltd.); low-speed refrigerated centrifuge(Eppendorf 5810R, Germany), ultra-low temperature freezer(Thermo Scientific -80°C, USA), optical microscope(Olympus BX53, Japan), electronic balance(Mettler-Toledo PL2002, Switzerland), pathological slide machine(Leica RM2245, Germany).

## 4 Animal husbandry conditions

All rats were housed in the SPF-grade animal laboratory of Henan Academy of Traditional Chinese Medicine^[18]^. The ambient temperature was controlled at 22±2°C,the relative humidity was 50%-60%, and the circadian rhythm was 12h light/12h darkness^[19]^. They were given free access to standard rat pellet feed(Beijing Huafukang Biotechnology Co., Ltd.) and underwent acclimatization feeding for 48 hours^[20]^. During this period, the rats’ activity, diet, water intake, and defecation were observed daily^[21]^. The experiment was started after confirming that there were no abnormalities^[22]^.

## 5 Methods^[23]^

Ten SD rats were randomly divided into four groups using a random number table: the normal group(number 2, female) received no intervention and was fed a standard diet; the control group(number 8, male) received only saline enema treatment^[24]^; the TNBS group(numbers 1 and 7, males; numbers 3 and 4, females) used TNBS/ethanol mixture enema to establish the rat model; and the DSS group(numbers 9 and 10, males; numbers 5 and 6, females) used DSS free drinking water to establish the rat model^[25]^. All methods were carried out in accordance with relevant guidelines and regulations, including the ARRIVE guidelines and the AVMA Guidelines for the Euthanasia of Animals(2020 edition)^[17]^.

### 5.1 Anesthesia^[26]^

Before modeling, rats in the control and TNBS groups were anesthetized with isoflurane(5% for induction, 2-3% for maintenance) in an oxygen flow of 0.8 L/min using a small animal anesthesia machine^[27]^. Adequate anesthesia was confirmed by the absence of pedal withdrawal and corneal reflexes. This method was chosen in accordance with the AVMA Guidelines for the Euthanasia of Animals to minimize pain and distress^[17]^. After the corneal reflex disappeared and the limb muscles relaxed, an enema was performed to avoid intestinal damage caused by struggling.

### 5.2 TNBS modeling^[28]^

TNBS was dissolved in 50% ethanol to prepare a modeling solution of 100 mg/kg. A rat-specific gavage needle(10 cm length, 2 mm diameter) was lubricated with saline and gently inserted 8 cm proximal to the anus into the distal colon. The TNBS/ethanol solution was injected over 10-15 seconds, followed by 0.3 mL of air to clear the needle. The rat was held in a head-down vertical position for 2 minutes to prevent leakage. After the rats regained consciousness from anesthesia, they were returned to their cages and allowed to resume normal food and water intake.

### 5.3 DSS group modeling^[29]^

Freshly prepared 3% DSS solution was used to replace regular drinking water, and rats were allowed to drink freely for 7 days. During this period, the DSS solution was changed daily to ensure a stable solution concentration. After the modeling was completed, normal drinking water was restored.

### 5.4 Control group treatment^[30]^

The same volume of normal saline was administered as the TNBS group for enema. The procedure, anesthesia method and body position were kept the same as the TNBS group to eliminate the interference of the enema procedure itself on the experimental results. This procedure-matched design is a critical methodological feature that has been omitted in most prior TNBS/DSS comparison studies, and it allows attribution of colonic pathology specifically to the chemical reagent rather than procedural trauma.

## 6 Observation and detection indicators

### 6.1 General condition monitoring^[31]^

During the modeling period, the body weight of each group of rats was recorded at 9:00 am every day, and the characteristics of feces and blood in feces were observed and recorded.

### 6.2 Specimen collection and processing^[32-34]^

#### 6.2.1 Blood samples

After the model was established, all rats were fed for 12 hours and fasted for 12 hours. Whole blood was collected by puncturing the abdominal aorta using a 10ml intravenous infusion set and a syringe. The blood was aliquoted into 5ml centrifuge tubes and allowed to stand for 24 hours at 4°C. The supernatant was then transferred to a 2ml centrifuge tube and centrifuged at 3000rpm for 10 minutes at 4°C to separate the serum. The serum was then frozen at -80°C for ELISA detection.

#### 6.2.2 Colon tissue specimens

After the rats were euthanized by CO_2_inhalation(flow rate: 20% chamber volume/min) followed by cervical dislocation^[35]^, the entire colon from the anus to the cecum was quickly dissected^[17]^. The total length was measured by an electronic balance and photographed. The intestinal tract was cut along the mesenteric margin, rinsed with physiological saline, and the mucosal congestion, edema, erosion and ulceration were observed by the naked eye. The lesion area 2-8 cm away from the anus was selected and cut into 2 cm long tissue segments. One segment was fixed in 4% paraformaldehyde solution at 4°C. The remaining tissues were packaged and transported on crushed ice and frozen at -80°C for later use.

#### 6.2.3 Pathological and histological analysis

After the fixed colon tissue was dehydrated in a gradient, embedded in paraffin, sectioned at 5 μm, dewaxed to water, and stained with HE, the integrity of the intestinal mucosa structure, the arrangement of glands, the degree of inflammatory cell infiltration and the depth of ulcers were observed under an optical microscope. The modified Macpherson pathological scoring criteria were used to make quantitative scores(0-10 points, the higher the score, the more serious the damage).

#### 6.2.4 Serum cytokine detection

Serum TNF-α and IL-6 levels were detected by ELISA. The operation was strictly carried out in accordance with the kit instructions. The absorbance value at 450 nm was read by the microplate reader, and the sample concentration was calculated according to the standard curve.

## 7 Statistical methods

Data analysis was performed using SPSS 26.0 statistical software. Quantitative data were expressed as mean±standard deviation(x±s). One-way ANOVA was used for comparisons between groups, and LSD-t test was used for pairwise comparisons. χ^2^ test was used for categorical data. *P<*0.05 was considered statistically significant. Coefficient of variation(CV% = SD/mean × 100%) was calculated for each endpoint within model groups as a measure of intra-group variability. Exploratory sex-stratified data are presented descriptively without statistical inference due to limited per-sex sample sizes(n=2).

## 8 Results

### 8.1 Changes in general condition of rats

During the modeling period, rats in the normal and control groups showed good mental state, active behavior, normal food and water intake, continuous weight gain, and normal stool characteristics without bloody stools. The absence of inflammatory signs in the procedure-matched control group(saline enema, same anesthesia protocol) provides direct evidence that enema-associated mechanical trauma does not independently confound inflammatory readouts at the doses and technique used in this study. In the TNBS group, rats showed lethargy and reduced activity 24 hours after modeling, and significant diarrhea appeared 48 hours later, with loose or watery stools; three rats showed visible bloody stools. In the DSS group, rats began to show decreased food intake on day 3 of modeling, diarrhea on day 5, and bloody stools in some rats on day 6; the symptoms were milder than in the TNBS group.

### 8.2 Morphological changes of the colon

In both the normal and control groups, the colon was soft and elastic, with smooth, red mucosa, and no signs of congestion, edema, or erosion. In the TNBS group, the colon was stiff and dark in color, with obvious congestion, edema, scattered erosions, and ulcers on the mucosal surface. The ulcers were mostly concentrated in the distal colon and showed a focal distribution. In the DSS group, the damage was more diffuse, involving the entire colon, with larger ulcer areas but shallower depths(Figure 2).

**Figure 2:**
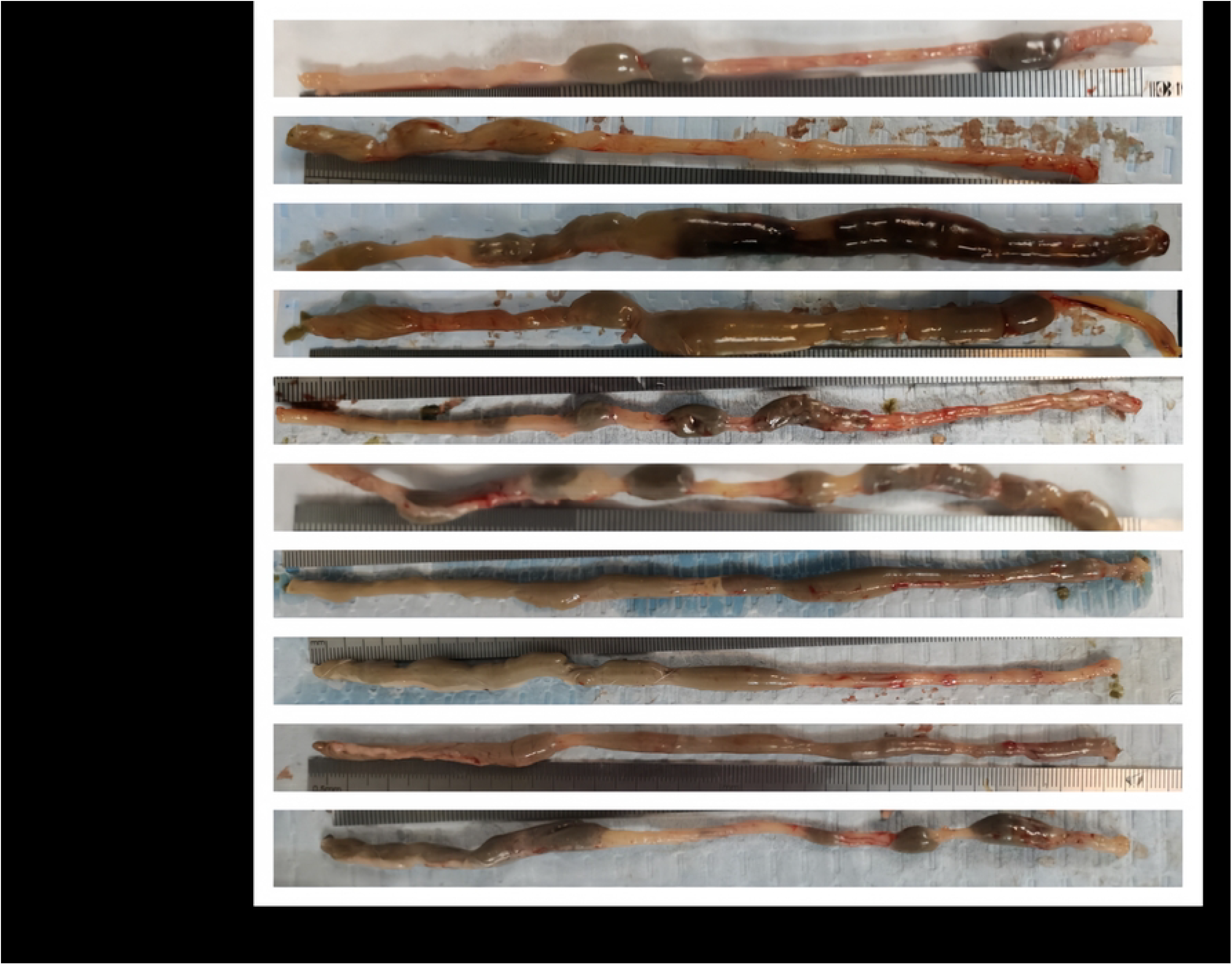

### 8.3 Pathological and histological results

HE staining showed that the intestinal mucosa structure was intact in both the normal and control groups, with glands arranged neatly and only a small number of lymphocytes infiltrating the lamina propria. In the TNBS group, the intestinal mucosa structure was severely damaged, with a large number of glands necrotic and sloughed off, and diffuse neutrophil infiltration in the lamina propria and submucosa, with ulcers extending to the submucosa. In the DSS group, the intestinal mucosa integrity was destroyed, with disordered gland arrangement and partial atrophy, and mixed infiltration of neutrophils and lymphocytes in the lamina propria. The ulcers were shallower and confined to the mucosal layer(Figure 3).

**Figure 3:**
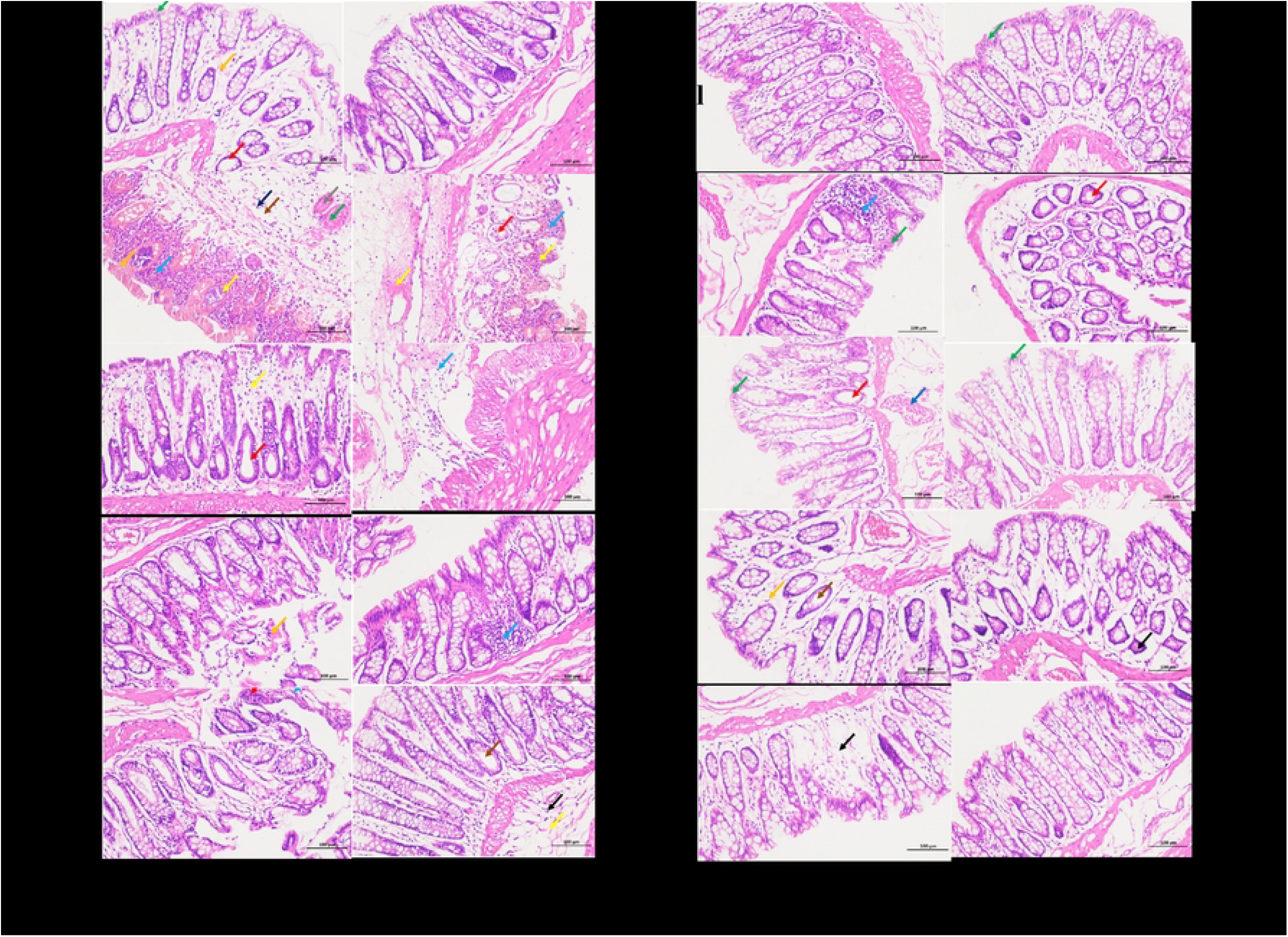

### 8.4 Serum cytokine levels

In the normal group and the control group, TNF-α levels were(24.8±3.2) pg/ml and(25.6±3.0) pg/ml, respectively, and the IL-6 levels were(18.2±2.6) pg/ml and(18.8±2.5) pg/ml, respectively, with no significant difference between the two groups(*P>*0.05), confirming no cytokine elevation from the enema procedure. In the TNBS group, serum TNF-α and IL-6 levels increased to(90.2±10.5) pg/ml and(66.5±8.3) pg/ml, respectively, while in the DSS group they were(73.5±9.8) pg/ml and(53.2±7.6) pg/ml, respectively. The cytokine levels in both groups were significantly higher than those in the normal group and the control group(*P<*0.05), with a greater increase in the TNBS group(*P<*0.05). Detailed serum TNF-α and IL-6 levels for each group are presented in Table 1.

**Table 1.** Serum TNF-α and IL-6 expression levels(pg/ml) in rats of each group 7 days after rat modeling.

| Group | Normal group | Control group | TNBS | DSS |
| --- | --- | --- | --- | --- |
| TNF-α | 24.8 ± 3.2 | 25.6 ± 3.0 | 90.2 ± 10.5 ▲ * | 73.5 ± 9.8 ▲ * △ |
| IL-6 | 18.2 ± 2.6 | 18.8 ± 2.5 | 66.5 ± 8.3 ▲ * | 53.2 ± 7.6 ▲ * △ |

Note: Compared with the normal group , $\blacktriangle P < 0.05$ ; compared with the control group , \* $P < 0.05$ ; compared with TNBS , $\triangle P < 0.05$ .

Post-hoc exploratory analysis: Coefficient of variation(CV%) for TNF-α was 11.6% in the TNBS group vs. 13.3% in the DSS group; for IL-6, 12.5% vs. 14.3%. Although both groups showed moderate variability at this sample size, the trend toward slightly lower CV in the TNBS group is consistent with the more rapid and stereotyped chemical injury of this model. Larger sample sizes are needed to confirm these differences. Sex-stratified exploratory data indicated that, in the TNBS group, female rats showed numerically higher TNF-α than males(mean difference approximately 8–12 pg/mL); in the DSS group, this sex difference was smaller. These observations are preliminary given n=2 per sex per group, but warrant investigation in future studies with larger, sex-balanced cohorts. All post-hoc analyses are strictly exploratory and should be interpreted accordingly.

## 9 Discussion

The pathological mechanism of ulcerative colitis(UC) is complex, with immune dysregulation being a core element^[36-38]^. The construction of animal models must closely mimic the pathological characteristics of human UC^[38, 39]^. The results of this study confirmed that both TNBS and DSS reliably induced UC-like pathology in rats, consistent with prior literature. However, the most important finding of this study is methodological rather than confirmatory: the procedure-matched saline enema control group demonstrated that enema-associated mechanical trauma did not independently elevate serum TNF-α, IL-6, or pathological scores, directly validating the attribution of all inflammatory readouts in the TNBS group to the chemical agent. This is a critical control that is systematically absent in the majority of published TNBS/DSS comparison studies.

TNBS enema and DSS free drinking water methods are currently the mainstream approaches for constructing UC models, and their modeling mechanisms differ fundamentally. TNBS, as a hapten, binds to colonic mucosal proteins to form a complete antigen, inducing a specific T-lymphocyte-mediated immune response with large-scale release of pro-inflammatory cytokines and acute focal mucosal damage; this model has a short modeling period and a severe inflammatory response, similar to the pathological characteristics of acute exacerbations of human UC. DSS, on the other hand, disrupts tight junctions between colonic mucosal epithelial cells, causing barrier failure and secondary innate immune activation^[40]^; its course is progressive, more closely resembling the chronic, recurrent clinical characteristics of human UC. This mechanistic distinction maps directly onto two immunologically distinct endotypes increasingly recognized in human UC research: a Th1/Th17-polarized endotype(characterized by high IFN-γ and TNF-α, associated with steroid-refractory disease) and an innate barrier-disruption endotype(characterized by high IL-1β and epithelial permeability markers). The significantly higher TNF-α and IL-6 in the TNBS group(*P<*0.05), combined with focal transmural lesion distribution, is consistent with Th1 endotype alignment; the diffuse mucosal-confined DSS lesions are consistent with innate barrier-disruption endotype alignment.

The results of this study showed that both modeling methods successfully induced a rat model of ulcerative colitis, exhibiting typical clinical symptoms such as weight loss, diarrhea, and hematochezia, shortened colon length, significant mucosal damage and inflammatory cell infiltration, and significantly elevated serum inflammatory factors(TNF-α and IL-6), consistent with previous findings. TNF-α, as a key pro-inflammatory factor, can induce apoptosis of intestinal epithelial cells and promote inflammatory cell infiltration; IL-6 participates in the amplification and persistence of the inflammatory response, and elevated levels of both are important markers of inflammatory activation in UC. The TNBS group showed a greater increase in inflammatory factors, suggesting a more severe inflammatory response consistent with T-lymphocyte-mediated immune injury; the DSS group had relatively lower cytokine levels, consistent with a less specific barrier-disruption mechanism. In terms of model characteristics, the TNBS group exhibits rapid onset of action, with obvious symptoms appearing within 24 hours. Pathological damage is primarily characterized by focal ulcers in the distal colon, making it suitable for studies of acute inflammation mechanisms, screening of T-cell-targeted therapies and potent anti-inflammatory drugs, and preliminary exploration of intestinal damage repair mechanisms. However, this model suffers from an overly severe inflammatory response and a short disease course, making it difficult to simulate the chronic, recurrent nature of human UC, and carries a higher risk of mortality. The DSS group, on the other hand, has a milder modeling process, with symptoms gradually worsening and diffuse damage distributed throughout the colon. Pathological changes are primarily mucosal inflammation, better reflecting the pathological characteristics of total colonic involvement and chronic inflammation in human UC. It is suitable for chronic disease observation, epithelial barrier repair research, microbiome-modulating drug development, and immunomodulatory drug development. Furthermore, the DSS modeling procedure is simple, requiring no anesthesia or enema, reducing the interference of operative trauma on experimental results and improving model reproducibility.

This study, by including a procedure-matched control group, definitively excluded damage to colonic tissue caused by the enema procedure itself, ensuring that all pathological changes in the TNBS group were attributable to the chemical reagent rather than mechanical trauma, thus establishing a more rigorous methodological standard for TNBS/DSS comparative research. However, this study also has certain limitations: the small sample size(n=2–4 per group) limits the statistical power and may affect the generalizability of findings, including the sex-stratified sub-analysis which is strictly exploratory; the lack of long-term follow-up prevented observation of relapse rates in the two models; and the detection of only two inflammatory factors, TNF-α and IL-6, is insufficient for a comprehensive analysis of the full inflammatory landscape. Future research should expand the sample size with balanced sex stratification, extend the observation period, and add detection indicators such as IL-1β, IL-10, IFN-γ, Th1/Th17 cell subsets, gut microbiota profiling, and epithelial permeability markers, to further validate the immune endotype framework and refine the comparative analysis of the two models.

The details of the modeling process are crucial to model stability: During TNBS enema, the insertion depth must be strictly controlled(8cm) to avoid spillage; maintaining an inverted position for 2 minutes ensures sufficient contact between the medication and the colonic mucosa. DSS solution must be freshly prepared and changed daily to prevent concentration reduction or bacterial contamination from affecting the modeling effect. Using an intravenous infusion set combined with a 10ml syringe during blood collection improves the success rate and avoids interference from hemolysis on serum test results. Standardizing these operational details is key to ensuring model reproducibility and can provide a reference for subsequent research.

## 10 Model-selection decision framework

Based on the findings of this study, the following three decision framework is proposed for selecting between TNBS and DSS models in preclinical UC research:

### 10.1 Research objective

Acute inflammation mechanism, T-cell biology, anti-TNF drug screening: TNBS; Chronic disease course, barrier repair, microbiome, mucosal healing, DSS.

### 10.2 Technical constraints

Isoflurane anesthesia equipment available and trained operator present: TNBS feasible; No anesthesia capacity or solo operator: DSS preferred(no anesthesia required, lower procedural variability).

### 10.3 Control group design

TNBS experiment, what must include procedure-matched saline enema control(anesthesia and enema, no TNBS) to isolate chemical from mechanical injury; DSS experiment, standard drinking-water control is sufficient.

## 11 Conclusion

This study makes three contributions beyond confirming that TNBS and DSS induce UC in rats. First, inclusion of a procedure-matched saline enema control group, systematically absent in most prior comparison studies, validates the attribution of inflammatory readouts to the chemical reagent rather than mechanical enema trauma, establishing a more rigorous methodological standard for TNBS/DSS comparative research. Second, the differential cytokine profiles(TNBS: higher TNF-α and IL-6, focal transmural lesions; DSS: lower cytokines, diffuse mucosal injury) are reinterpreted within an immune endotype framework: TNBS models Th1-polarized UC, while DSS models innate barrier-disruption UC, a distinction with direct implications for translating preclinical findings to human UC subtypes. Third, a mechanism-of-action-based three-step model-selection decision framework is provided, offering actionable guidance for preclinical drug screening that goes beyond the generic “acute vs. chronic” recommendations of prior comparison papers. Both the TNBS enema method and the DSS free drinking method can successfully establish a rat model of ulcerative colitis; researchers should choose the most suitable modeling method based on experimental objectives, research duration, and the mechanism of action of the investigational therapeutic, guided by the framework presented here. These contributions collectively establish a more rigorous and translationally relevant standard for animal model selection in UC research.

## Acknowledgements

This work was supported by the Third Batch of Henan Province’s Traditional Chinese Medicine Young Talent Training Program of China and the joint research project funded by the National Center for Inheritance and Innovation of Traditional Chinese Medicine of China(Grant No. 2024ZXZX1102).

## Ethical Considerations

This work was supported by the Animal Ethics Committee of Henan Academy of Traditional Chinese Medicine(Ethics approval number: 20230911).

## Author contributions

Yongkang An: Funding acquisition, Writing-review & editing

Fang Wu: Investigation, Writing-original draft

Linhan Zhang: Formal analysis

Menghua Shi: Investigation

Shuangxi Zhang: Methodology, Supervision, Writing – review & editing

Xiang’an Zhang: Supervision, Writing – review & editing

All authors approved the final manuscript.

## Data availability

We confirm that any datasets generated during and/or analyzed during the current study are publicly available, available upon reasonable request, and data sharing is applicable to this article.

